# Genome-wide Association Study of Clinical Features in the Schizophrenia Psychiatric Genomics Consortium: Confirmation of Polygenic Effect on Negative Symptoms

**DOI:** 10.1101/161349

**Authors:** Tim B. Bigdeli, Roseann E. Peterson, Stephan Ripke, Silviu-Alin Bacanu, Richard L. Amdur, Pablo V. Gejman, Douglas F. Levinson, Brien P. Riley, David St. Clair, Marcella Rietschel, James T.R. Walters, Roel A. Ophoff, Andrew McQuillin, Hugh Gurling, Dan Rujescu, Patrick F. Sullivan, George Kirov, Michele T. Pato, Carlos N. Pato, Ole A. Andreassen, Michael J. Owen, Michael C. O'Donovan, Aiden Corvin, Anil K Malhotra, Bryan J. Mowry, Tõnu Esko, Thomas Werge, Kenneth S. Kendler, Schizophrenia Working Group of the Psychiatric Genomics Consortium, Ayman H. Fanous

## Abstract

Schizophrenia is a clinically heterogeneous disorder. Proposed revisions in *DSM - 5* included dimensional measurement of different symptom domains. We sought to identify common genetic variants influencing these dimensions, and confirm a previous association between polygenic risk of schizophrenia and the severity of negative symptoms. The Psychiatric Genomics Consortium study of schizophrenia comprised 8,432 cases of European ancestry with available clinical phenotype data. Symptoms averaged over the course of illness were assessed using the *OPCRIT, PANSS, LDPS, SCAN, SCID, and CASH*. Factor analyses of each constituent *PGC* study identified positive, negative, manic, and depressive symptom dimensions. We examined the relationship between the resultant symptom dimensions and aggregate polygenic risk scores indexing risk of schizophrenia. We performed genome - wide association study (*GWAS*) of each quantitative traits using linear regression and adjusting for significant effects of sex and ancestry. The negative symptom factor was significantly associated with polygene risk scores for schizophrenia, confirming a previous, suggestive finding by our group in a smaller sample, though explaining only a small fraction of the variance. In subsequent *GWAS*, we observed the strongest evidence of association for the positive and negative symptom factors, with *SNPs* in *RFX8* on 2q11.2 (P = 6.27×10^-8^) and upstream of *WDR72 / UNC13C* on 15q21.3 (*P* = 7.59×10^-8^), respectively. We report evidence of association of novel modifier loci for schizophrenia, though no single locus attained established genome - wide significance criteria. As this may have been due to insufficient statistical power, follow - up in additional samples is warranted. Importantly, we replicated our previous finding that polygenic risk explains at least some of the variance in negative symptoms, a core illness dimension.

## Introduction

The symptomatic presentation, course, outcome, and severity of schizophrenia (SZ) have long been known to vary widely from patient to patient. *DSM* and *ICD* subtypes have their historical antecedents in the historical rubrics paranoia, hebephrenia, and catatonia, all of which predate the modern concept of SZ ^1^. Whether such putative, categorical subtypes best describe variation in clinical features is unclear, as many patients meet criteria for the undifferentiated subtype^2^. Dimensional constructs such as positive, negative, disorganization, and mood symptoms have been proposed using both theoretical and empirical criteria^3^, and evidence from family and twin studies suggests that these are heritable ^4^. Of major interest to researchers and clinicians alike is the potential of symptomatic dimensions for predicting outcome, severity, and treatment response.

We have previously suggested that genes could affect clinical variation in at least two ways ^4^. Modifier genes might influence the clinical character of the illness without increasing disease susceptibility. The most elegant examples of modifier genes per se are found in studies of cystic fibrosis. Though caused by mutations in the *CFTR* gene, the associated clinical features of cystic fibrosis, some of which influence mortality, are influenced by genes having no contribution to the risk of illness ^5^. Genetic modifiers of disease severity have been identified for a range of diseases, from facioscapulohumeral muscular dystrophy to Alzheimer’s disease ^6,7^. Second, allelic variation in disease susceptibility genes themselves might influence clinical characteristics, or predispose to a particular illness subtype. Such subtype - specific loci, or “susceptibility - modifier genes”, have been identified in several disorders, including Huntington’s disease, breast cancer and frontotemporal lobar degeneration ^8-11^.

Identification of both classes of genes in SZ could be beneficial in several ways. First, this might yield clues to underlying pathophysiological processes more or less specific to a clinical subtype or symptom dimension. This has the potential to inform questions of diagnosis and classification such as the use of dimensional approaches, as was debated in the development of DSM - 5 ^12^ but ultimately not adopted. Intriguingly, the demonstration of pleiotropic effects — both of specific loci and in the form of shared polygenetic liability — may hold additional clues. A recent study found that a polygenic risk score for bipolar disorder significantly predicted the severity of manic symptoms in SZ ^13^. Also noteworthy are reported copy - number variant (*CNV*) associations shared between SZ, autism and intellectual disability ^14,15^, which suggests a link between negative symptom - like psychopathology and the potentially debilitating effects of gene - disrupting CNVs ^16,17^. Importantly, such insights could provide leads in the development of novel treatments, since it is symptom dimensions, rather than the illness itself, that are ameliorated by medications.

We previously published a large genome - wide association study of positive, negative, and mood symptoms in a sample of 2,454 cases, which provided suggestive evidence of novel modifier genes influencing these dimensions ^18^. Furthermore, the polygenic risk of SZ predicted negative / disorganized symptom severity. In this report, we have used a similar approach in over 8,000 cases from the Psychiatric Genomics Consortium (PGC) study of SZ ^19^. Firstly, we sought to replicate our previous observation that polygenic risk of SZ influences the severity of negative symptoms. We also hypothesized that, with enhanced statistical power, we might detect additional evidence of modifier loci influencing classic clinical dimensions of psychotic illness.

## Methods

### Ascertainment and Assessment

The subsamples included in this study comprise 19 constituent sites from Stage 2 of the Psychiatric Genomics Consortium study of SZ. Ascertainment, diagnostic assessment, genotyping, and genotype quality control have been previously described ^19,20^. Briefly, 52 samples from the US, Europe, and Australia comprising 34,241 cases, 45,604 controls and 1,235 parent affected - offspring trios were genotyped using a number of commercial single nucleotide polymorphism (SNP) genotyping platforms. These data were processed using the stringent PGC quality control procedures, followed by imputation of SNPs and insertion - deletions using the 1000 Genomes Project reference panel (UCSC hg19/NCBI 37) ^21^ using IMPUTE2 ^22^, resulting in nearly *9.5M* markers for GWAS analysis.

In the current study, we focus on clinical dimensions of illness as the phenotypes of interest, and have performed a case - only GWAS of these quantitative traits. Clinical dimensions were derived directly from the individual items of symptom checklists or rating scales assessed in the individual PGC sites if they were available. These instruments included the Comprehensive Assessment of Symptoms and History (CASH) ^23^, Lifetime Dimensions of SZ Scale (LDPS) ^24^, Operational Criteria Checklist for Psychotic Illness (OPCRIT) ^25^, Positive and Negative Syndrome Scale (PANSS) ^26^. For some subsamples, we used the individual items of the diagnostic interviews, including the Schedule for Clinical Assessment in Neuropsychiatry (SCAN) ^27^ and Structured Clinical Interview for DSM - III - R (SCID) ^28^.The individual sites, their sample sizes, and the instruments used for developing quantitative phenotypes are presented in Supplemental Table 1.

**Table 1.** Association of PGC2 - trained polygenic scores with symptom dimensions of SZ. For each symptom dimension, the total N of cases, ***P*** - value threshold (P_T_) yielding the greatest predictive value, variance explained by this score (R^2^), and its significance (***P*** - value) are given for the full sample (i.e., “all PGC2”), the MGS study alone, and all non - MGS PGC2 studies.

| Symptom | Cohort | $N$ | $P_T$ | $R^2$ | $P$-value |
| --- | --- | --- | --- | --- | --- |
| Positive | MGS only | 2 386 | 0.1 | $1.53 \times 10^{-3}$ | 0.0558 |
| | non-MGS PGC2 | 5 944 | $10^{-4}$ | $1.07 \times 10^{-4}$ | 0.422 |
| | all PGC2 | 8 330 | 0.1 | $6.46 \times 10^{-5}$ | 0.214 |
| Negative | MGS only | 2 386 | 0.1 | 0.00452 | $9.95 \times 10^{-4}$ |
| | non-MGS PGC2 | 6 041 | 0.1 | 0.00373 | $1.83 \times 10^{-6}$ |
| | all PGC2 | 8 427 | 0.1 | 0.00386 | $6.39 \times 10^{-9}$ |
| Manic | MGS only | 2 384 | $10^{-4}$ | $3.50 \times 10^{-3}$ | 0.00375 |
| | non-MGS PGC2 | 4 566 | 0.2 | $9.49 \times 10^{-5}$ | 0.511 |
| | all PGC2 | 6 950 | $10^{-4}$ | $6.39 \times 10^{-4}$ | 0.0204 |
| Depressive | MGS only | 2 386 | 0.1 | $1.27 \times 10^{-3}$ | 0.0783 |
| | non-MGS PGC2 | 4 578 | 0.001 | $4.43 \times 10^{-4}$ | 0.153 |
| | all PGC2 | 6 964 | 0.1 | $3.84 \times 10^{-4}$ | 0.0546 |

### Quantitative phenotypes

Quantitative traits for the Stage 1 sites were derived from the individual items of assessment instruments as previously described ^13^. In brief, exploratory factor analysis of each individual site’s data, followed by confirmatory factor analysis, identified four factors that were most prominent across sites: Positive, Negative, Manic, and Depressive symptoms. We then harmonized these factors across sites and instruments, then Z - transformed factor scores within each site. For the current expansion into the Stage 2 sites, we opted to retain the four - factor model. We selected the relevant items comprising these factors from the instruments used in the Stage 2 sites, and used them to create factors scores, which were subsequently Z - transformed. A more detailed description of these methods is given in the accompanying Supplemental Note.

### Polygenic scoring analyses

To test for a polygenic effect on symptom dimensions, we performed risk score profiling as previously described ^19,29^. Briefly, SZ risk scores were generated for each studcts included in the training set, while offering improved power and less sample attrition compared to subdividing the full cohort into approximate halves. We computed several scores based on varying *P* - value threshold signifying the proportion of SNPs with smaller *P* - values in the training set; *P* - value thresholds ranged between 0.0001 and 1.0. We tested for association between symptom factors and SZ polygenic scores by linear regression, adjusting for study - site and all covariates used in the primary association analyses. To investigate possible heterogeneity among study sites, we also analyzed each site separately and combined site - specific results by fixed - effects meta - analysis.

### Genome - wide association and meta - analysis

We tested for association between SNPs and each trait by linear regression, as implemented in PLINK v1.07 (http://pngu.mgh.harvard.edu/~purcell/plink/) ^30^, using allelic dosages and adjusting for significant covariates including sex and ancestry principal components (where applicable). For each trait, relevant covariates were identified for the full cohort by backwards - stepwise regression, after adjusting for study site. We performed GWAS of each trait separately for individual study sites, combining summary statistics in subsequent random - effects meta - analyses using *METAL* ^31^. We excluded all SNPs with minor allele frequency (*MAF*) less than 0.01, average statistical imputation information (INFO) less than 0.5, or absent from more than half of total number of sub - studies.

For X chromosome SNPs, tests of association were carried out separately for male and female subjects. For each study site, we meta - analyzed these male - and female - specific results, subsequently combining the resultant summary statistics across study sites in a second round of meta - analysis.

### Gene - set enrichment analysis

We applied a new method, Data - Driven Expression Prioritized Integration for Complex Traits (*DEPICT*) ^32^, to identify significantly enriched gene – sets / pathways and tissues / cell - types. Briefly, genes harboring or in the vicinity of associated SNPs are tested for enrichment for “reconstituted” gene sets, comprised of curated sets expanded to include coregulated loci. Tissue and cell type enrichment analysis is conducted by testing whether genes were highly expressed in any of 209 *MeSH* annotations based on microarray data for the Affymetrix U133 Plus 2.0 Array platform. As input, we used SNPs significant at the recommended *P* - value inclusion thresholds of 5×10^-8^ and 10^-5^; we relaxed the latter threshold to 5×10^-5^ to allow for the inclusion of a greater number of associated loci.

## Results

### Exploratory factor analysis

As previously described, we used a combination of scree plots and clinical judgment to select a four - factor model consisting of positive, negative, manic, and depressive symptoms ^13^. We observed that these four symptom dimensions were the most salient in the EFAs of the seven OPCRIT sites. EFA of the PANSS in three sites (Oslo, Munich, and CATIE) revealed separate negative and disorganized factors, as did a published EFA of the CASH in the one site using that instrument in this study. In our previous EFA of the LDPS in the MGS sites, there was a single negative symptom factor which also included disorganization symptoms. We therefore used a single negative factor, and combined negative and disorganization symptoms into one factor in the PANSS and CASH sites.

We observed separate manic and depressive factors in the OPCRIT, CASH and PANSS sites, but not the LDPS sites. We therefore opted to divide the LDPS mood factor into separate manic and depressive factors. As different instruments have differences in emphasis and granularity, we sought to test whether they tapped into underlying constructs that were more or less the same. In the Dublin sample, the Pearson product - moment correlations for the negative, manic, and depressive factors across the PANSS and OPCRIT were, respectively, 0.85, 0.70, and 0.48 (Supplementary table S7).

### Polygenic scoring analyses

Following replicated evidence for a polygenic contribution to SZ ^29^, we assessed whether weighted polygenic risk scores based on a training - set of cases and controls ^19^ could predict each symptom factor in an independent target dataset, namely each constituent dataset included herein. Given the requirement that training and target datasets be independent, scores were generated via a “leave - one - out” procedure (see Methods). Results for scores based on varying P - value thresholds are displayed in Figure 1a. Overall, we observe the strongest relationship with polygenic risk for the negative symptom factor. We observed the most significant such enrichment for a score constructed of SNPs with P - values less than 0.1 in the PGC2 GWAS of SZ (*R*^2^ = 0.00386; *P* = 6.39×10^-9^). We observed little or no evidence of association with polygenic risk for the manic (*R*^2^ = 6.39×10^-4^; *P* = 0.0204), depressive (*R*^2^ = 3.84×10^-4^; *P* = 0.0546), and positive symptom factors (*R*^2^ = 6.46×10^-5^; *P* = 0.214).

**Figure 1:**
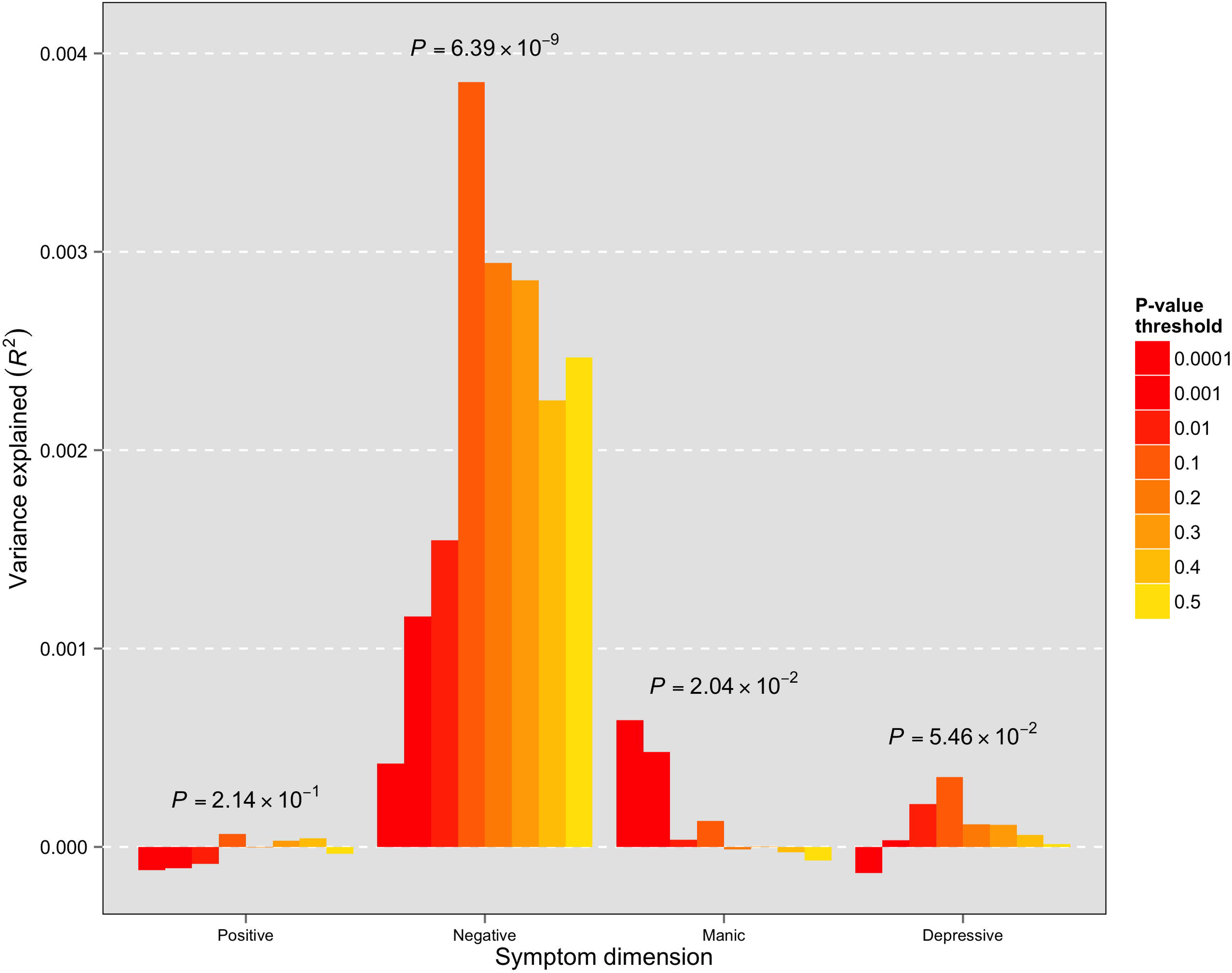
(a) Predictive value of SCZ polygene scores. Results based on varying SNP ***P*** - value inclusion thresholds are grouped by symptom dimension along the x - axis. Proportion of variance explained (adjusted R^2^) is shown on the y - axis. Displayed ***P*** - values correspond to the inclusion thresholds yielding the largest proportion of variance explained. (b) Forest plot displaying study - specific results for the association of SCZ polygene scores with negative symptoms. pT is the selected ***P*** - value threshold for the poygenic score tested; for each site, beta, se, p, and n are the estimated effect and its standard error from linear regression, the corresponding ***P*** - value, and the number of cases with available clinical data, respectively.

Given a previously observed association between SZ risk and negative symptom severity in the MGS study ^18^, we sought to assess to what extent this particular study might be driving the observed pattern of findings. In Table 1, we report the significance and predictive value of the SZ polygenic score as applied to each symptom dimension in the full PGC sample, the MGS study alone, and excluding MGS from the combined PGC sample. For negative symptoms, exclusion of the MGS study yielded nearly identical results with respect to variance explained (*R*^2^ = 0.0037), though with an attendant diminution of statistical significance resulting from the smaller sample size (P = 1.83×10^-6^). Examination of the other symptom dimensions reveals a lesser degree of consistency across cohorts.

We followed - up the possibility of heterogeneity across study sites by individually analyzing each based on the overall best *P* - value threshold and subsequently combining study - specific results by fixed - effects meta - analysis. For negative symptoms, a forest plot displaying estimated effects for each site is given in Figure1b; we did not observe significant heterogeneity in effects sizes across sites (Cochran’s *P* = 0.0955). Corresponding forest plots for depressive, manic, and positive symptoms are included in the Supplemental material; of these, only positive symptoms demonstrated significant heterogeneity across study sites (Cochran’s *P* = 0.0003).

### Genome - wide association and meta - analysis

Genomic inflation factors (λ) were 0.993, 1.014, 1.0, 1.003 for the positive, negative, manic, and depressive factors, respectively. We observed the strongest evidence of association overall for the positive symptom factor on 2q11.2 at SNPs in *RFX8* (*P* = 6.27×10^-8^), followed by the negative symptom factor on 15q21.3, with SNPs upstream of *WDR72* (*P* = 7.59×10^-8^). Regional association and forest plots for these two SNPs are provided in the accompanying supplemental information. Although no single SNP surpassed established genome - wide significance criteria (i.e., 5×10^-8^), we observed several moderate associations (P < 10^-6^) at distinct chromosomal loci (Figure 2); 5, 4, and 3 for the positive, negative, and depressive symptom factors, respectively. These are summarized, with respect to the two strongest single SNP associations, in Table 2.

**Table 2.** Association results for SNPs attaining P < 10^-6^ in primary GWAS of symptom dimensions. For each SNP, Chr and Mb give its genomic coordinates (hg19); INFO is the statistical imputation information; Freq is the frequency of the reference (first listed) allele, and Z - score is its estimated effect; P and PSZ are the ***P*** - values in the current and full PGC case - control analyses, respectively. The nearest gene within 1Mb is shown; its position relative to a gene is given parenthetically (negative and positive kb values indicate up - and downstream positions).

| Symptom | Chr | Mb | SNP | Alleles | INFO | Freq | Z | P | P<sub>SZ</sub> | Nearest Gene (kb) |
| --- | --- | --- | --- | --- | --- | --- | --- | --- | --- | --- |
| Positive | 1p13.3 | 107.5 | rs7522504 | C/G | 0.981 | 0.327 | 4.96 | $7.24 \times 10^{-7}$ | 0.802 | *PRMT6* (-102.2kb) |
| | 2p24.2 | 19.08 | rs72778999 | A/T | 0.726 | 0.919 | 4.93 | $8.35 \times 10^{-7}$ | 0.0874 | *NT5C1B* (+306kb) |
| | 2q11.2 | 102.08 | rs74263251 | A/C | 0.913 | 0.934 | 5.41 | $6.27 \times 10^{-8}$ | 0.973 | *RFX8* (0) |
| | 8q21.12 | 80.02 | rs78234477 | C/T | 0.951 | 0.014 | -5.12 | $3.06 \times 10^{-7}$ | 0.878 | *IL7* (+304.4kb) |
| | 22q12.1 | 27.61 | rs474758 | C/T | 0.977 | 0.781 | 4.91 | $8.94 \times 10^{-7}$ | 0.195 | *LOC101929539* (+294.4kb) |
| Negative | 6q16.2 | 99.99 | rs56238691 | C/T | 0.805 | 0.014 | 5.19 | $2.15 \times 10^{-7}$ | 0.904 | *CCNC* (0) |
| | 14q22.3 | 57.93 | rs2484490 | C/T | 0.917 | 0.522 | 4.9 | $9.44 \times 10^{-7}$ | 0.0636 | *C14orf105* (-5.61kb) |
| | 15q21.3 | 54.12 | rs144529410 | C/G | 0.717 | 0.026 | 5.38 | $7.59 \times 10^{-8}$ | 0.531 | *UNC13C* (-188.3kb) |
| | 16q23.1 | 77.85 | rs3909915 | C/G | 0.748 | 0.985 | -5.13 | $2.85 \times 10^{-7}$ | 0.890 | *VAT1L* (0) |
| Depressive | 2q33.3 | 206.1 | rs145452099 | C/T | 0.694 | 0.097 | 4.96 | $7.09 \times 10^{-7}$ | 0.149 | *PARD3B* (0) |
| | 3p26.1 | 8.31 | rs59441182 | A/G | 0.954 | 0.64 | 5.05 | $4.33 \times 10^{-7}$ | 0.391 | *LMCD1-AS1* (0) |
| | 3q25.2 | 153.95 | rs112126680 | G/T | 0.688 | 0.963 | 4.96 | $6.96 \times 10^{-7}$ | 0.224 | *ARHGEF26* (0) |

**Figure 2:**
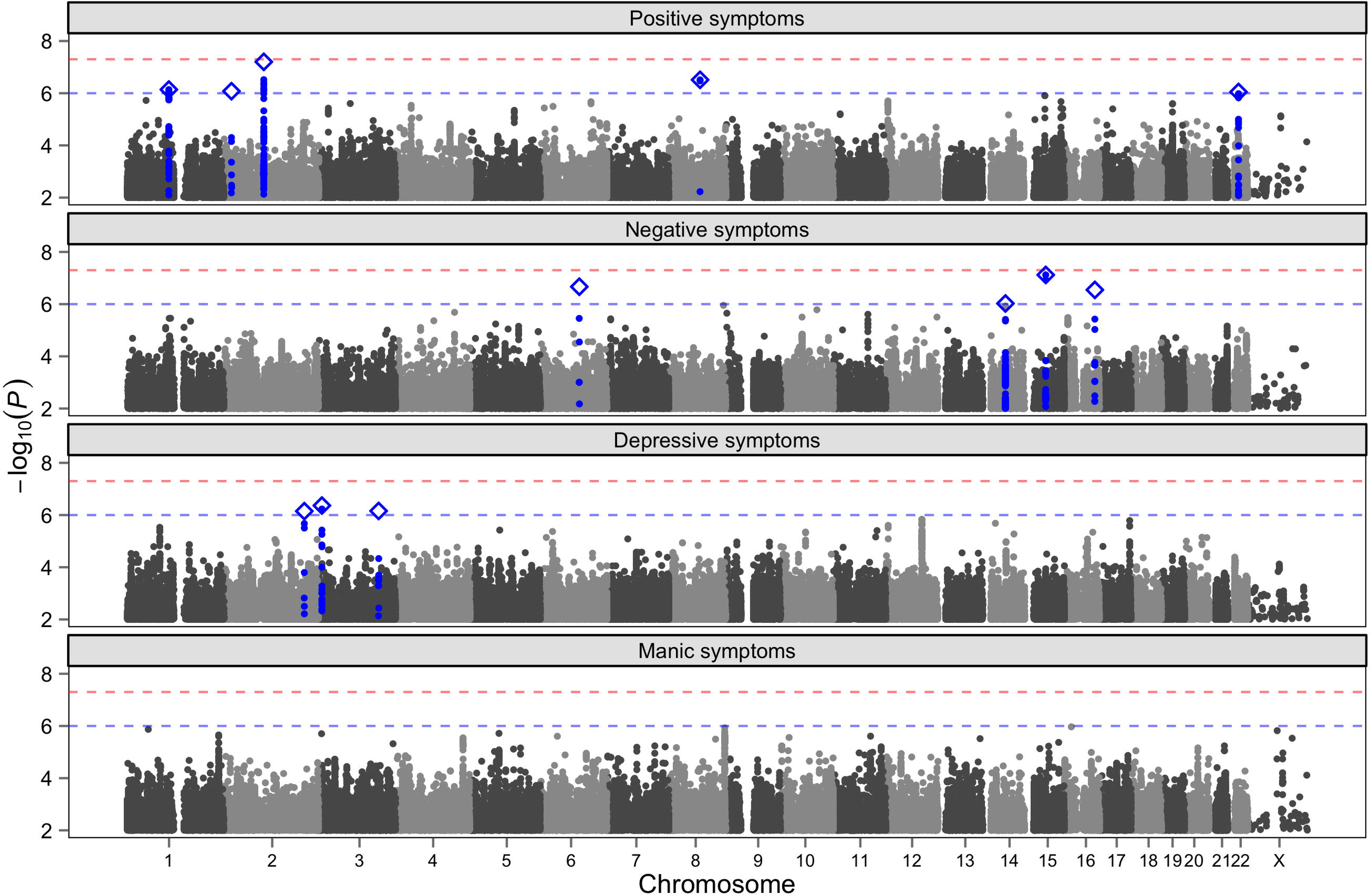
Manhattan plots for GWAS of four clinical dimensions. Red and blue horizontal lines show thresholds for genome - wide (5×10^-8^) and suggestive (10^-6^) significance, respectively. For suggestively associated SNPs, the most significant “independent” marker within a 500kb region is represented as a blue diamond; neighboring SNPs in linkage disequilibrium (r^2^ > 0.1) with this most significant marker are highlighted.

### Gene - set enrichment analyses

We investigated whether particular gene - sets were enriched for associations with negative symptoms using DEPICT. For SNPs significant at *P* < 10^-5^, representing 29 independent loci, a single pathway, response to drug (GO:0042493), was significantly enriched (FDR < 0.2), or contained more significant genes than expected by chance, after correction for multiple testing. This gene - ontology (GO) category is defined as any process that results in a change of state or activity of a cell or an organism in response to a drug stimulus, and includes both response to antipsychotic drug (GO:0097332) and response to antidepressant (GO:0036276) as child terms (14).

We also considered a more inclusive threshold of *P* < 5×10^-5^, which yielded a total of 109 independent loci. A total of seven (7) gene - sets were found to be significantly enriched following correction for multiple testing. Of particular note is the observed enrichment of protein - protein - interaction (PPI) networks of low - density lipoprotein - related proteins 1 and 2 (FDR < 0.01), and ankyrin - 2 (FDR < 0.05).

For each SNP *P* - value threshold considered, Table 3 gives the gene - sets that remained significant after correction for multiple testing (FDR < 0.20).

**Table 3.** Gene - set enrichment analysis results for negative symptoms. Nominal ***P*** - values and estimated false - discovery rate (FDR) for suggestive gene - sets (FDR < 0.20), based on analyses of SNPs significant at a specified ***P*** - value threshold. For each SNP ***P*** - value threshold considered, the total number of independent loci considered (pairwise r2 < 0.1 within 500kb) is given parenthetically.

| *P*-value threshold<br>(No. loci) | Gene-set ID | Description | *P*-value | FDR |
| --- | --- | --- | --- | --- |
| $1 \times 10^{-5}$ (29) | GO:0042493 | response to drug | $3.85 \times 10^{-7}$ | < 0.05 |
| $5 \times 10^{-5}$ (109) | ENSG00000081479 | LRP2 PPI subnetwork | $1.81 \times 10^{-6}$ | < 0.01 |
| | ENSG00000123384 | LRP1 PPI subnetwork | $5.47 \times 10^{-6}$ | < 0.01 |
| | MP:0004056 | abnormal myocardium compact layer morphology | $2.40 \times 10^{-5}$ | < 0.05 |
| | ENSG00000145362 | ANK2 PPI subnetwork | $3.24 \times 10^{-5}$ | < 0.05 |
| | ENSG00000188612 | SUMO2 PPI subnetwork | $7.34 \times 10^{-5}$ | < 0.20 |
| | GO:0042417 | dopamine metabolic process | $1.27 \times 10^{-4}$ | < 0.20 |
| | GO:0010657 | muscle cell apoptotic process | $1.72 \times 10^{-4}$ | < 0.20 |

## Discussion

To our knowledge, this is the largest GWAS of SZ - related quantitative traits performed to date. Of particular interest is our finding of a significant correlation between polygenic risk of SZ and severity of negative symptoms. This replicates, in a much larger multi - national sample, a previous observation of this relationship in the MGS sample ^18^, and to a greater degree of statistical significance. More broadly, this finding provides additional support for genetic factors underlying the observed clinical heterogeneity in SZ^4^, and that at least some risk genes for SZ are in fact susceptibility - modifier genes. Some of these genes and their gene products might therefore provide clues to developing more effective treatments for this often - disabling symptom dimension.

Experiment - wide, the strongest evidence of single - marker association was observed for positive symptoms, at SNPs in ***RFX Family Member 8, Lacking RFX DNA Binding Domain (RFX8)***, which encodes a potential transcription factor but which has not previously been implicated in SZ. An association between intergenic SNPs at 1p13.3 and negative symptoms also approached, albeit without attaining, genome - wide significance (*P* < 10^-7^). These SNPs fall between, and upstream of *WD repeat domain 72 (WDR72)* and unc - 13 homolog C (UNC13C). In particular, *UNC13C* encodes a protein essential to presynaptic transmission and synaptic plasticity ^33,34^, and has been implicated in several neuropsychiatric traits, including post - traumatic stress disorder ^35,36^.

At more modest significance thresholds (*P* < 10^-6^), we observed several associations between the positive, negative, and depressive symptoms factors and SNPs mapping to both genes and intergenic regions, which vary with respect to the degrees of support for a role in SZ etiology. Notably, these SNPs were not significantly associated with SZ risk (P > 0.05) in the full PGC study ^19^. While our single SNP results could be due to a paucity of true allelic effects on these clinical dimensions of illness, in our view, this is more likely due to insufficient power to detect small allelic effects in even a large sample such as this. This is suggested by the limited success in identifying replicable genome - wide significant effects in sample sizes comparable to ours or larger, in studies of heritable quantitative traits such as personality ^37,38^,. The GWAS literature in general suggests that in some complex traits, very large studies and considerable phenotypic refinement are required to detect small effect genes ^39,40^.

That exploratory gene - set enrichment analyses highlighted several etiologically relevant molecular networks would also seem to support this interpretation. The observed enrichment of **LRP1** and **ANK2** protein - protein interaction networks for associations with negative symptoms is particularly salient, given the implication of the former in Alzheimer’s disease ^41^ and, more recently, of the ***ANK2*** gene in autism ^42,43^, which have features of the cognitive and social aspects of negative symptoms.

Perhaps most convincing, however, is the highly significant association between negative symptom severity and polygenic risk of SZ, suggesting that many SZ risk loci also influence its clinical presentation. That SZ risk was only found to predict negative symptom severity may reflect the temporal stability of negative symptoms and their more accurate assessment by interviewers, thus contributing to a higher signal - to - noise ratio compared to the other dimensions ^1,44,45^. However, the finding that polygenic risk of bipolar disorder significantly predicts the less reliably assessed manic symptom factor in SZ ^13^ suggests that this not the case. It is possible that negative symptoms are a marker for cases of SZ that arise more directly from common variants or the rare variants that are in LD with them, while phenocopies are more likely to be related to environmental etiologies, such as substance misuse. This is supported by the observation that cocaine - and amphetamine - induced psychotic states tend to present with more positive than negative symptoms ^46^. The differential impact of the polygenic risks of bipolar disorder (Ruderfer et al., 2014) versus SZ (as reported here) suggests potential biological underpinnings of both the commonalities and differences between these two disorders, whose relationship has been the subject of considerable debate ^47-49^. For example, bipolar disorder polygenes might explain some of their observed phenomenological overlap, in the form of the manic - like symptoms seen in both, while SZ polygenes might explain phenomena more specific to SZ, such as negative symptoms.

The heterogeneity of the assessment instruments could have increased the false negative rate by decreasing the signal - to - noise ratio in GWAS. The instruments used in this study differ in purpose, content, and granularity. The **SCID** ^28^ and **SCAN** ^27^ are structured interviews designed to make diagnoses rather than comprehensively inventory signs and symptoms. The **OPCRIT** ^25^, while it is a comprehensive checklist of classic signs and symptoms of SZ and affective disorders, is also meant to aid in diagnosis using multiple criteria. The **PANSS** ^26^, on the other hand, is not intended for diagnosis, but rather, for a fine - grained assessment of positive and negative symptoms of SZ, for purposes such as determining treatment efficacy. Unlike diagnostic interviews, it is therefore lacking in items reflecting classic manic symptoms, and instead, more sensitively assesses psychotic excitement and agitation. Similarly, **PANSS** depressive items do not comprehensively include classic neurovegetative symptoms, perhaps resulting in the lower **PANSS - OPCRIT** correlation we observed for the negative factor compared to others.

Because the PANSS is designed to assess common positive and negative symptoms of **SZ, PANSS** factors tended to be more normally distributed. The **OPCRIT’s** depressive and especially, manic factors demonstrated considerable positive skew, in part because of the relatively low prevalence of classic manic symptoms, as compared to excitement and hostility, in SZ. Although the “manic” constructs in the **OPCRIT** and **PANSS** are therefore different, it was reassuring that they were strongly correlated in the Dublin sample, suggesting that they index the same underlying construct. Nevertheless, their divergent distributions and item content could have resulted in false negative results when analyzed together.

Related to the heterogeneity of the instruments used is the heterogeneity in factor solutions across sites. In order to maximize statistical power, we opted to use the most prominent and compelling set of clinical dimensions across sites, in all of the sites. This resulted in our using the less than best - fitting models in a some of the sites, which could have reduced power in these sites, and concomitantly, in the study as a whole. Our combining of two factors into one in some of the sites (PANSS, SCAN, SCID) might have diluted a putative genetic signal affecting one of these factors alone, but not the other one. These limitations highlight the need for standardized ratings in future quantitative measures of symptoms.

An additional limitation of this study was the lack of inclusion of potentially important covariates, including environmental factors such as cannabis use and urbanicity, as well as cognitive impairment. The latter in particular might be related to negative and disorganization symptoms. However, environmental factors occurring prior to the onset of illness are difficult to reliably determine retrospectively, and were therefore largely unavailable in these samples.

An important next step will be to attempt replication of these findings in additional independent datasets, including newer additions to the Psychiatric Genomics Consortium. This could include samples of patients with bipolar disorder and/or major depressive disorder, who sometimes manifest the some of the same symptoms seen in SZ. Furthermore, as large numbers of efficacy studies have used the **PANSS**, it might be possible to expand future studies such as ours with more routine biobanking and genotyping in such studies.

## Acknowledgements

The work specific to this report was funded by the United States Department of Veterans Affairs Merit Review Program (5I01CX000278) to A.H.F. This project has received funding from the European Union’s Seventh Framework Programme for research, technological development and demonstration under grant agreement no 279227 (M.R.).

Statistical analyses were carried out on the Genetic Cluster Computer (http://www.geneticcluster.org) which is financially supported by the Netherlands Scientific Organization (NWO 480-05-003) along with a supplement from the Dutch Brain Foundation and the VU University Amsterdam.

